# DEEPCON: Protein Contact Prediction using Dilated Convolutional Neural Networks with Dropout

**DOI:** 10.1101/590455

**Authors:** Badri Adhikari

**Affiliations:** Department of Mathematics and Computer Science, University of Missouri-St. Louis, St. Louis, MO 63121, USA

## Abstract

**Background:** Exciting new opportunities have arisen to solve the protein contact prediction problem from the progress in neural networks and the availability of a large number of homologous sequences through high-throughput sequencing. In this work, we study how deep convolutional neural network methods (ConvNets) may be best designed and developed to solve this long-standing problem.

**Method:** With publicly available datasets, we designed and trained various ConvNet architectures. We tested several recent deep learning techniques including wide residual networks, dropouts, and dilated convolutions. We studied the improvements in the precision of medium-range and long-range contacts, and compared the performance of our best architectures with the ones used in existing state-of-the-art methods.

**Results:** The proposed ConvNet architectures predict contacts with significantly more precision than the architectures used in several state-of-the-art methods. When trained using the DeepCov dataset consisting of 3,456 proteins and tested on PSICOV dataset of 150 proteins, our architectures achieve up to 15% higher precision when L/2 long-range contacts are evaluated. Similarly, when trained using the DNCON2 dataset consisting of 1,426 proteins and tested on 84 protein domains in the CASP12 dataset, our single network achieves 4.8% higher precision than the ensembled DNCON2 method when top L long-range contacts are evaluated. DEEPCON will be made publicly available at https://github.com/badriadhikari/DEEPCON/.

## 1. Introduction

For a protein whose amino acid sequence is obtained using a protein-sequencing device, three-dimensional (3D) models may be predicted using template modeling or ab-initio. Template-modeling methods search for other similar protein sequences in a database of sequences whose 3D structures have already been experimentally determined using wet-lab experiments and use them to predict 3D models of the input sequence. The total number of protein structures determined through experimental methods such as X-ray crystallography and Nuclear magnetic resonance spectroscopy are currently limited to 147,817 as of January 2019 [1]. Protein sequences for which such templates cannot be found need to be predicted ab-initio, i.e. without the use of any structural templates. For structure prediction of protein sequences whose structural templates are not found, predicted protein contacts serve as the driver for folding [2,3].

Residue-residue contacts or inter-residue contacts (or just contacts) define which pairs of amino acids should be close to each other in the 3D structure, i.e. pairs that are in contact should remain close and those that are not should stay as far as possible. As defined in the Critical Assessment of Protein Structure Prediction (CASP) experiments [4,5], a pair of residues in a protein are defined to be in contact if their carbon beta atoms (carbon alpha for glycine) are closer than 8 Angstroms in the native (experimental) structure. In a true (or a predicted) contact matrix not all contacts are equally important. Local contacts, those with sequence separation less than 6 residues, and short-range contacts (with sequence separation between 6 and 12 residues) are not very useful for building an accurate 3D model. They are required for reconstructing local secondary structures but are not helpful to build folded proteins. However, medium-range contacts, the contact pairs that have sequence separation between 12 and 23 residues, and long-range contacts, the ones separated by at least 24 residues in the protein sequence, are necessary for building accurate models.

All top groups participating in the most recent Critical Assessment of Protein Structure Prediction (CASP) 13 experiment, including DeepMind’s AlphaFold method use contacts (or distance bins) for ab-initio protein structure prediction. These contacts can be predicted with relatively high precision for protein sequences that have hundreds to thousands of matches in protein sequence databases such as UNICLUST30 [6] and Uniref100 [7]. The sequence hits obtained, in the form of multiple sequence alignments (MSA) serve as an input to algorithms and machine learning methods to predict contact maps. While the overall goal in protein structure prediction is to predict three-dimensional models (3D information) from protein sequences (1D information), predicted protein contacts serve as the intermediate step (2D information). In the absence of machine learning, contacts are predicted from protein sequence alignments based on the principle that evolutionary pressures place constraints on the sequence evolution over generations [cite evfold]. The predicted contacts from these coevolution-based methods are a key input to machine-learning based methods that generally predict more accurate contacts.

The most successful contact prediction methods use convolutional neural networks (CNNs) fed with a combination of features generated from multiple sequence alignments and other sequence-derived features. After Jinbo Xu’s group first applied CNNs to predict contacts [3], CNNs have been found to be particularly well suited and highly effective for the contact prediction problem, mainly because of their ability to learn cross-channel (cross-feature) information (for example, the relationship between predicted solvent accessibility and predicted secondary structure). In [8] authors demonstrate that a basic CNN-based method can easily outperform another state-of-the-art meta-method based on basic neural networks. Similarly, in [9] authors demonstrate that a single CNN-based network delivers a remarkably better performance compared to a boosted deep belief network. Although the recent progress in contact prediction was initially attributed mostly to coevolutionary features generated using methods such as CCMpred [cite] and FreeContact [cite], recent findings [8,10] suggest that end-to-end training is possible in the near future, where the deep learning algorithm may contribute entirely to the performance and these hand engineered features may be found redundant. Most of the recently successful methods for contact predictions, as demonstrated by the recent CASP results, are available for public use. For instance, methods such as RaptorX [3], MetaPSICOV [11], DNCON2 [9], PconsC3 [12], and PconsC4 [2] are either available as a downloadable tool or a web server. Each of these methods use very different CNN architectures, different sets of input features, and self-curated datasets to train and benchmark their methods.

From the perspective of input and output data format, the protein contact prediction problem is similar to depth prediction [13] in computer vision, i.e. predicting 3D depth from 2D images. In the depth prediction problem, the input is an image of dimensions H × W × C, where H is height, W is width, and C is number of input channels, and output is a two-dimensional matrix of size H × W whose values represent the depth intensities. Similarly, in the protein contact prediction problem, the output is a contact probability map (matrix) of size L × L, and input is protein features of dimension L × L × N, where L is the length of the protein sequence and N is the number of input channels. Depth prediction usually involves three channels (red, green, and blue or hue, saturation, and value) while in the latter we have much higher number of features such as 56 or 441 [8]. Because of large number of input channels, the overall input volume becomes large, limiting the depth and width of deep learning architectures for training and testing. This also greatly affects the training time and requires high-end GPUs for training. These challenges, imposed by large number of input channels, are also observed in other problems such as plant genotype prediction from hyperspectral images. A protein contact matrix and its input features are symmetrical along the diagonal. All current methods (including this work) consider both upper and lower triangles (above and below the diagonal line) for training but only one of the triangles for evaluation.

The protein structure prediction problem has some additional unique characteristics. First, the input features for proteins are not all two dimensional. Input features such as length of a protein sequence are scalar or 0-dimensional (0D). For such input features, we create a channel with the same value throughout the channel. Similarly, input features such as secondary structure predictions are one-dimensional (1D) and we create two channels for each 1D input feature of length L - first channel by copying the vector L times to create a L × L matrix, and second channel by transposing the input feature vector and then copying L times to create a L × L matrix. Another way to generate 2D matrix from 1D vector is to compute outer product of the vector with its own transpose. Other input features which are 2D, are copied into the input volume, as they are. Also, the length of a protein sequence can vary from a few residues to a few thousand residues, implying a variable input feature volume. The dimension of each 2D feature (transformed into channels) will depend on the length of the protein. Unlike images protein structures cannot be scaled. Unlike real-world images, these input features, cannot be studied/understood visually towards understanding what the networks are learning. The training and test dataset that we can use is also limited. Unlike other publicly available datasets, the protein structure dataset size cannot be significantly increased because only up to around 11 thousand new structures are experimentally determined each year. What further reduces this dataset is the similarity between many of the structures deposited. On these proteins, if we perform some basic redundancy reductions such as keeping only the proteins with more than 20% sequence similarity and those with high resolution structures, the number of proteins available for training reduces to only around a five thousand [14]. It is yet to be explored how data augmentation methods may be applied to best exploit the data that is already available. Despite the fact that the data appears to be limited compared to many other datasets, some argue that it is sufficient to capture the principles of protein folding. In addition, a typical protein contact map, which is a binary matrix, is around 95% zeros and 5% ones [15].

The three-dimensional models of real-world objects and protein structures have a fundamental difference. Protein 3D models are not scalable. An object, such as a chair, in the real world may be tiny or large. Proteins can also be large or small but the size of structure patterns are always physically fixed. For instance, the size of an alpha helix (a common building block of many protein structures) is the same for proteins of any size. The distance between carbon-alpha atoms is a fixed physical distance whether the helices are in a small helical protein such as the mouse C-MYB protein or a large helical protein such as hemoglobin. Because of this unique characteristic of protein structures, it is yet to be fully understood how useful deep architectures such as U-Nets [16] can be for protein contact prediction although some groups have developed contact prediction methods using such architectures [2]. In addition, because of this rigid characteristic of protein structures (unlike images) it is not yet understood how data augmentation techniques for images such as cropping, scaling, rotation, and flipping can be used on these datasets. Similar to the problem of depth prediction, the ultimate goal in the field is to develop methods that can be scaled to predict physical distances map (in Angstroms). Since raw distance prediction is too difficult, some groups have developed methods that can predict contacts at various sequence separation thresholds [17] and demonstrate that such binning of distance ranges actually improves contact prediction at a single standard distance threshold, such as the standard of 8 Angstroms.

In this work, we demonstrate that dilated residual networks with dropout layers are best suited for addressing the protein contact prediction problem. It can be argued that the results in a small dataset may not hold true in large datasets. In addition, results generated using one kind of features may not hold true for another kinds of features. For instance, methods that work well for sequence-based features (such as secondary structures) may not work well for features generated from multiple-sequence alignments (such as covariance). Here, we experiment with two diverse datasets that use different input features. When trained on the DeepCov dataset consisting of 3,456 proteins, using the same dataset for training and testing our method achieves up to 6% and 15% higher precision on the PSICOV150 protein dataset when top L/5 and L/2 long-range contacts are evaluated, respectively (L is protein length). Similarly, when trained on the DNCON2 dataset consisting of 1,426 proteins, using the same dataset for training and testing, a single network in our method achieves 4.8% higher long-range precision (top L). While DNCON2 uses a two-stage approach combined with ensembling of 20 models, we use a single network. Although we trim the input length of a protein to 256 for all proteins in our training experiments, after the training, our models can make predictions for a protein of any length.

## 2. Materials and Methods

### Datasets

In our experiments we use two datasets: 1) the DNCON2 dataset [9] available at http://sysbio.rnet.missouri.edu/dncon2/ and 2) DeepCov dataset [8] available at https://github.com/psipred/DeepCov. Following the deep learning practice, for each experiment, we consider three subsets – training subset, validation subset, and an independent test set. The DNCON2 dataset consists of 1,426 proteins between 30 and 300 residues in length. The dataset was curated before the CASP10 experiment in 2012 and the protein structures in the dataset are of 0-2 Angstroms in resolution. These proteins are filtered by 30% sequence identity to remove redundancy. Of the full dataset, we use 1,230 proteins for training and the remaining 196 as the validation set. We found that the protein with PDB ID 2PNE in the validation set was skipped during evaluation in the DNCON2 method and hence we too skipped. We train and validate our method using the 1,426 proteins and test on 62 protein targets in the CASP12 experiment. The CASP12 protein targets range from 75 to 670 residues in length. For evaluating our method on the CASP12 dataset, we predict contacts for the whole protein target and evaluate the contacts on the full target (or domains). The DeepCov dataset consists of 3,456 proteins used to train and validate models, i.e. tune hyperparameters. The models trained using the DeepCov dataset are tested on the independent PSICOV dataset consisting of 150 proteins. In **Figure 1**, we summarize our two experimental setups.

**Figure 1.**
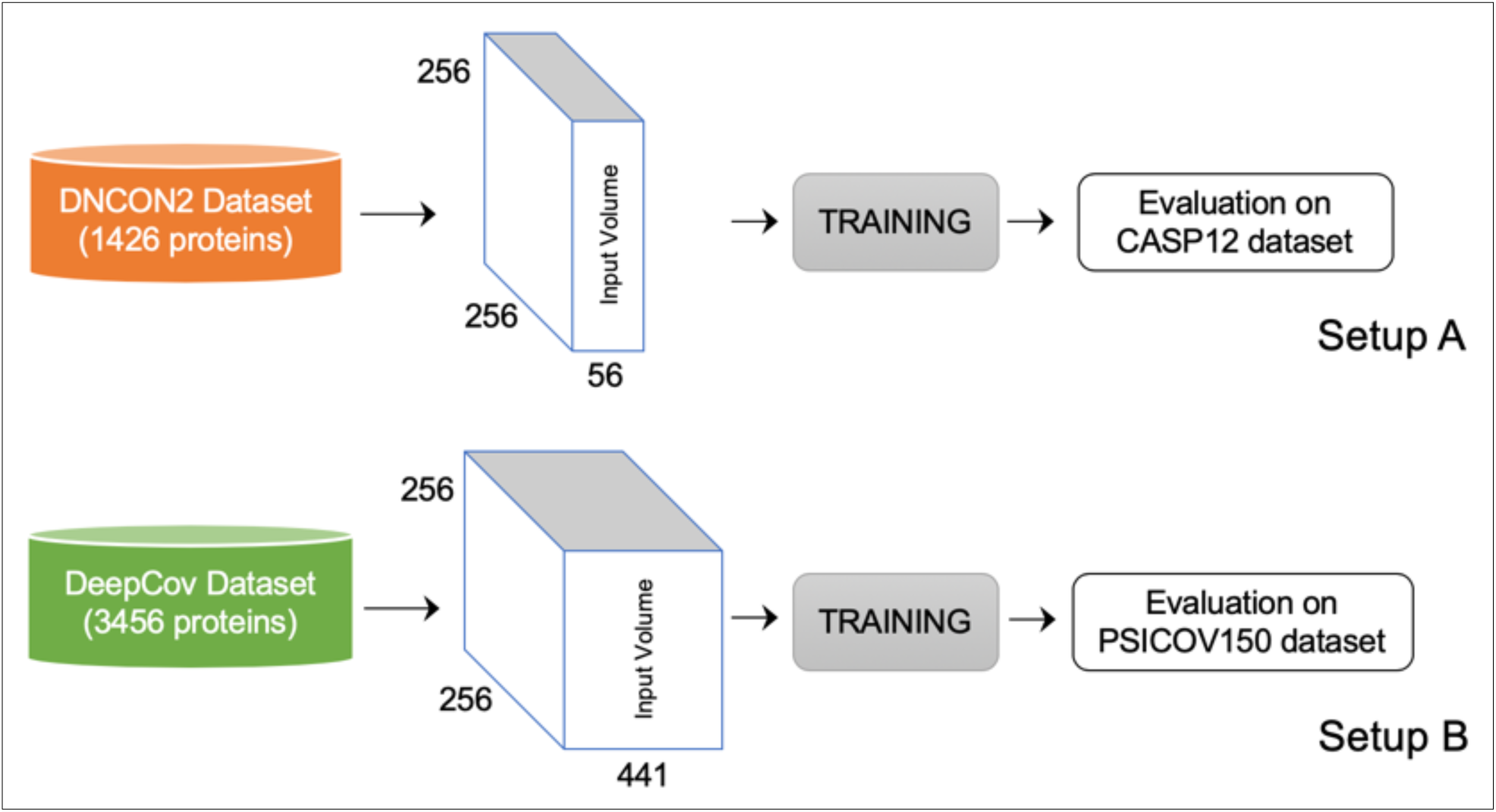
Setups designed for our experiments.

### Contact Evaluation

Following the standard in the field [4], as evaluation metrics of contact prediction accuracy, we use the precision of top L/5 predicted long-range contacts (P_TOP-L/5-LR_), where L is the length of the protein sequence. For a more rigorous approach, we evaluate our methods using the precision of top L/2 long-range contacts (P_TOP-L/2-LR_), precision of top L long-range contacts (P_TOP-L-LR_), and the precision of all medium- and long-range contacts (P_ALL-MLR_). For instance, to calculate P_ALL-MLR_ of contacts predicted for a protein with N_LR_ number of medium- and long-range contacts, we first round the top N_LR_ medium- and long-range predicted probabilities (after ranking the probabilities) to 1s and the rest to 0s. Precision is then calculated as the ratio of ‘the number of matches between predicted and true matrix’ and N_LR_. We find that evaluating all (not just some top) medium- and long-range contacts is both a more robust and effective evaluation metric for model selection.

### Input Features

In training experiments associated with DNCON2 dataset, we use eight groups of features as input for training and testing our models. These include scalar values such as sequence length, one-dimensional features such as solvent accessibility and secondary structure predictions, and two-dimensional features such as features computed from position specific scoring matrix, sequence separation information, pre-computed statistical potentials, features computed from input multiple sequence alignment, and coevolution predictions from the alignment. For generating multiple sequence alignments, we follow the approach of using HHsearch [18] followed by JackHmmer [19] as discussed in the DNCON2 method (see our DNCON2 paper for details). In total, we use 29 unique features as input. For each protein, we have 56 input channels in total. For each 0D (scalar) feature we copy the value into the entire channel. Similarly, we copy each 2D input feature as-is into the final input volume as a single channel. For each 1D input feature for a protein of length L, we create two channels - first channel by copying the vector L times to create a L × L matrix, and second channel by transposing the input feature vector and then copying L times to create an L × L matrix. Although the maximum length of a protein in this dataset is 300, we trim all input features to 256 length so that the longest protein has an input volume is 256 × 256 × 56. For training, all proteins with length less than 256 are 0-padded so that all input volumes have the same dimension of 256 × 256 × 56. For our experiments involving the DeepCov dataset, we use the 441 covariance features (channels) calculated using the publicly available script (cov21stat) and dataset in the DeepCov package. For this dataset the input volume for each protein is L × L × 441.

### Training Convolutional Neural Networks

Our ConvNets involve an input layer, a number of 2D convolutional layers with batch normalization or dropouts, residual connections and Rectified Linear Units (ReLU) activations. In all architectures, the final layer is a convolutional layer with one filter of size 3×3 followed by a ‘sigmoid’ activation to predict contact probabilities. Since we only have convolutional layers, the variables are the numbers and size of filters at each layer, and dilation rate when dilated convolutional layers are used. **Figure 2** summarizes our approach. All the CNN filters in the first layer convolve through the input volume of 256 × 256 × N producing batch normalized and ‘relu’ activated outputs passed as input to the subsequent layers. The number of channels, N, is 56 when DNCON2 dataset is used and 441 with DeepCov dataset is used. We stop training the model if the validation accuracy does not improve for 20 epochs and reduce the learning rate by 0.5 when the loss does not improve for 10 epochs. Error is computed using binary cross entropy calculated as −(y log(p)+(1−y) log(1−p)), where p is the output of the sigmoid activation of the last layer for each residue pair, and y is 1 if the residue pair are in contact in the experimental structure or else is 0. Although we crop the input length of our training proteins to 256, after the training the model can make predictions for a protein of any length. Since we do not use max pooling or any dense connections, a model of any arbitrary dimension can be built for predicting proteins of lengths longer than 256 residues. For instance, to predict contacts for a protein with length 500 (i.e. input volume of 500 × 500 × 56), we first build a model of the same input dimensions (i.e. 500 × 500) and then load all the trained weights into this new model and make predictions for the input volume. Since contact matrix is symmetrical, we average the prediction of either triangle to generate final predictions. We do not do any model ensembling.

**Figure 2.**
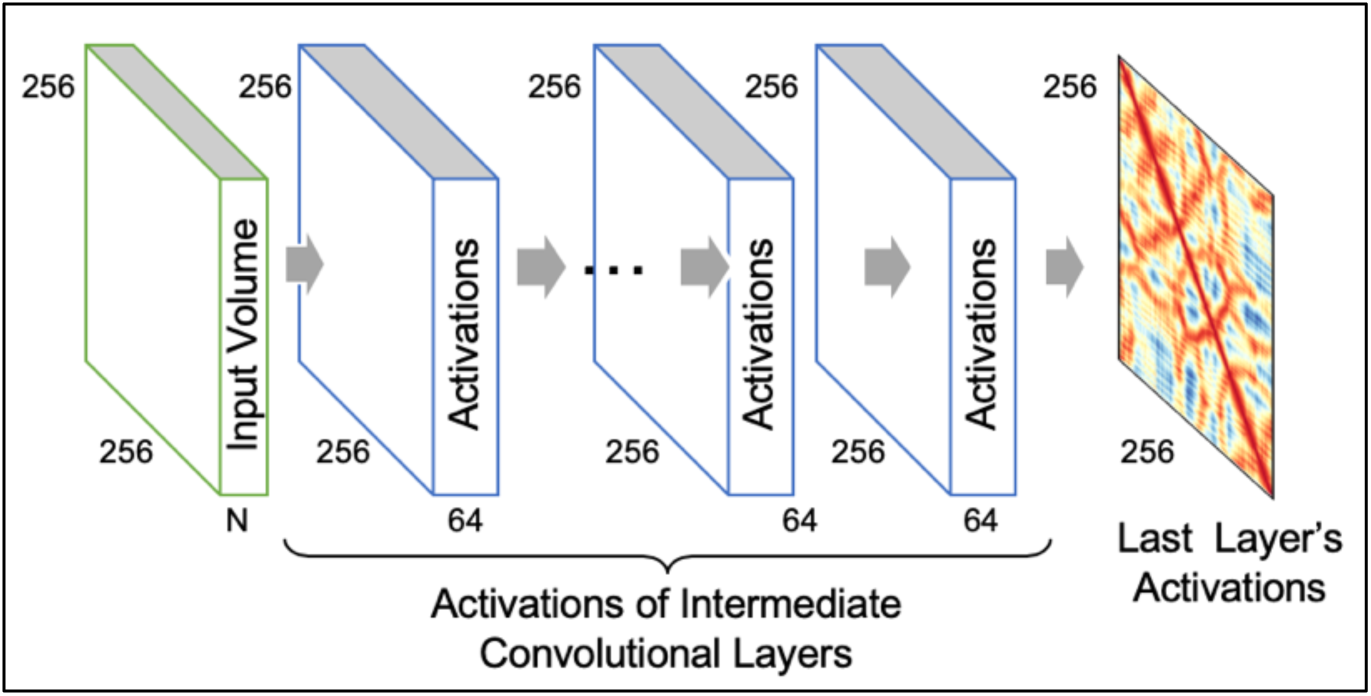
General architecture for training and testing. Filters in the first layer of convolutional neural network slide through the input volume for each protein with a 300×300×56 matrix (56 input channels) producing activation volumes. One filter in the last layer slides through the activation volume of the previous layer to generate the final activations at each position of the 256 × 256 matrix.

We used the Keras library (https://keras.io/) with Tensorflow (https://www.tensorflow.org/) backend for our method development. On a machine with 2 CPUs and 24 GB memory with a NVIDIA Tesla K80 GPU the training (around 35 to 45 epochs) takes about 16 hours for either of the DNCON2 or DeepCov dataset. For testing larger architectures we used the Tesla P100, V100, and Quadro P6000 GPUs.

### Network Architectures

We start our training experiments with standard convolutional neural networks (or fully convolutional networks), each preceded by a batch normalization layer and ReLU activation. We find that performance of such networks drop after 32 to 64 convolutional layers. Next, we design a residual block consisting of two convolutional layers, each preceded by a batch normalization layer and ReLU activation. Fixing the total number of convolutional filters in each layer to 64, we design four residual networks: 1) a regular residual network with depths (number of blocks) as the key parameter, 2) an architecture with the second batch normalization layer in each residual block replaced with a dropout layer, 3) we replace the last convolutional layer with a dilated convolutional layer with dilation rate alternating between 1, 2, and 4, and 4) an architecture that is a combination of the second and third architecture, i.e. we replace the second batch normalization layer with a dropout layer, and replace the second convolutional layer with a dilated convolution layer at alternating dilation rates of 1, 2, and 4. We also tested many other architecture variations such as filter sizes of 16, 24, and 32, and different ways to alternate between batch normalization layers and dropout layers. On average, these architectures were not significantly better than the four architectures we discuss here.

Using the DNCON2 dataset as our training and validation set and CASP12 targets as test dataset, we compared the precision of a fully convolutional network (FCN), and the four residual architectures – standard residual network, residual network with dilation, residual network with dropout, and residual network with dilation and dropout. On the fully convolutional networks and residual networks, we experimented adding dropout layers in many ways and found that alternating between batch normalization layers and dropout layers yield the best performance[20][20][20][20][20][20][20][20][20]. On our standard residual network with 64 layers having 64 3×3 filters in each layer, we tested dropout values of 0.2, 0.3, 0.4, and 0.5. When we evaluate these models on the precision of top L/5 long-range precision (P_TOP-L/5-LR_) and all medium-, and long-range precision (P_ALL-MLR_) we find that specific value of dropout does not matter as long as we have the dropout layers in place. For most of our experiments we fixed 0.3 as the parameter for our dropout layers (i.e. keep 70% weights). Since our findings resonate with the findings in [20] we also hypothesize that the issue of “diminishing feature reuse” is pronounced in the problem of contact prediction and that these dropout layers partially overcome the issue. In fact, we observed up to 6% gain in P_TOP-L/5-LR_ when we randomly replace some of the initial batch normalization layers in a fully convolutional layers with just 16 layers (each having 64 filters). These findings suggest that dropout layers, when used appropriately, can be highly effective in network architectures for protein contact prediction. When evaluated on the independent CASP12 dataset consisting of 62 targets, we find that residual networks with dilation and dropout outperform all the four other architectures, across all the precision metrics, P_TOP-L/5-LR_, P_TOP-L/2-LR_, P_TOP-L-LR_ and P_ALL-MLR_ (see **Figure 4**). We call this residual network with dilation and dropout as the DEEPCON method.

**Figure 3.**
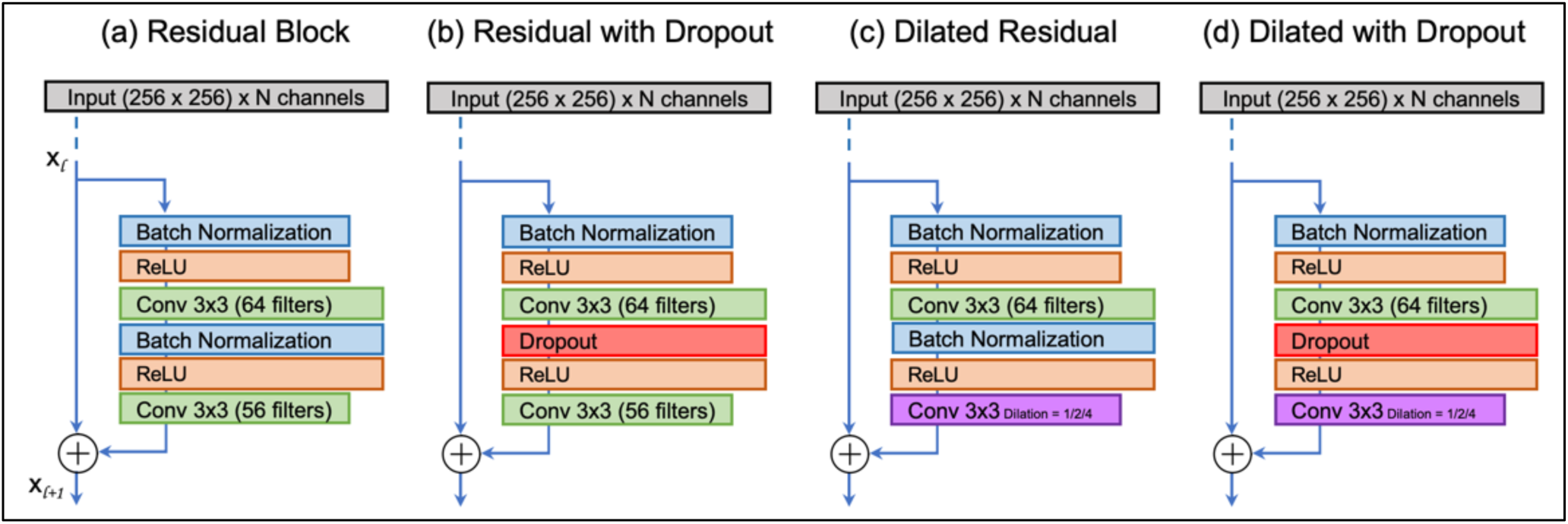
Residual network architectures designed for training and evaluation.

**Figure 4.**
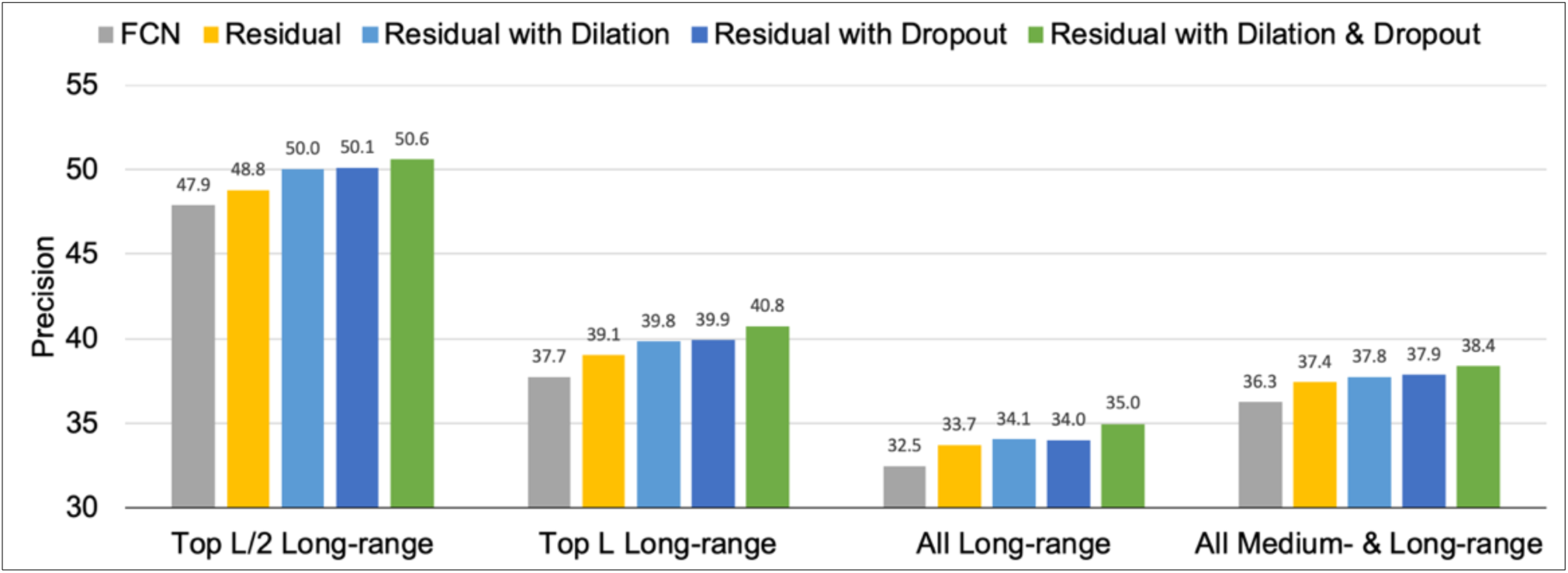
Comparison of the precision on CASP12 dataset obtained from various architectures when models are trained and validated using the DNCON2 dataset.

## 3. Results

### Comparison of Network Architectures with Other Network Architectures

To benchmark our DEEPCON architecture we compare it with the architectures used in the current state-of-the-art methods – Raptor-X [3], DNCON2 [9], DeepCov [8], and PconsC4 [2]. The Raptor-X method uses residual networks with around 60 convolutional layers. The DNCON2 method uses 6 convolutional layers each with 16 filters and an additional convolutional layer with one filter for generating the final contact probabilities. This network has around 50 thousand parameters. In the DeepCov method, 441 input channels are reduced to 64 using a Maxout layer, and then fully convolutional layers of various depths are tested. DeepCov performs best at the receptive field size of 15, i.e. 7 convolutional layers. The PconsC4 method directly uses the U-Net architecture that accepts 256 × 256 × 64 input volume, i.e. input channels will be first projected to 64 channels. Such an architecture has 31 million parameters. In our setup A, where we use DeepCov dataset for training and validation and PSICOV 150 proteins as the test dataset, we find that DEEPCON performs similar to the standard residual architecture (see **Figure 5)** when the evaluation metric is P_TOP-L/5-LR_. On the P_ALL-LR_ and P_ALL-MLR_ metrics, however, DEEPCON’s performance is significantly higher than the standard residual-type network architecture, a fully connected CNN architecture, and the U-Net architecture. In our setup B, where we use DNCON2 dataset for training and CASP12 protein domains for testing, we observe similar results (see **Figure 5**). Compared to the DeepCov dataset, the DNCON2 dataset consists of much lesser input features and smaller number of training proteins. Hence, we repeated our training for each architecture two times to obtain a more accurate comparison. To validate our findings, we designed a third setup - we generated DNCON2 like features for the 3,456 proteins in the DeepCov dataset and tested the performance on the PSICOV150 proteins. In this setup observed DEEPCON and the top ranking method. It is worth noting that that these two datasets (DNCON2 and DeepCov) have highly dissimilar input features. While DNCON2 uses features such as secondary structure predictions as input, DeepCov uses covariance calculations entirely from the multiple sequence alignments. DEEPCON’s high performance on these two very different datasets suggests that deep residual networks with dropout are reliable and work well for a variety of input features towards predicting protein contacts.

**Figure 5.**
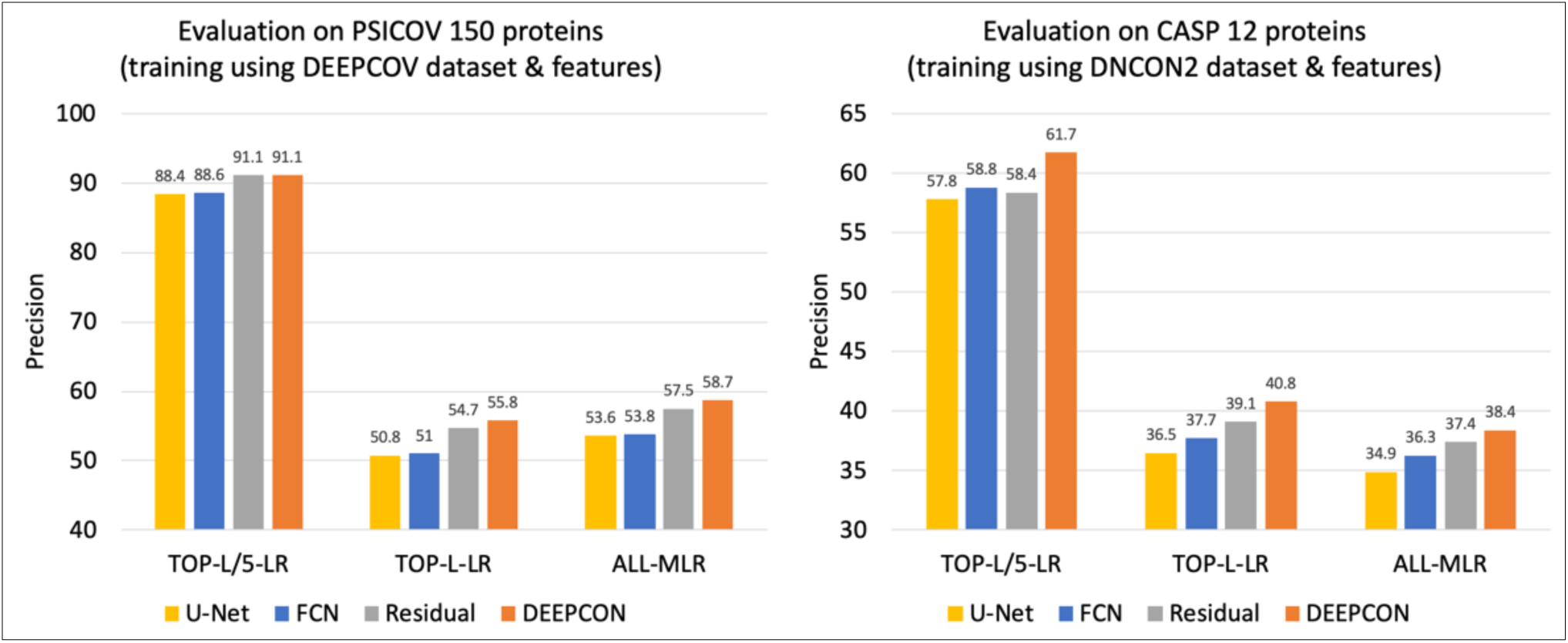
Comparison of the performance of various ConvNet architectures – fully connected ConvNets (FCN), residual networks with dropout and dilation (DEEPCON), regular residual network, and U-Net like architecture – using P_TOP-L/5-LR_, P_TOP-L-LR,_ and P_ALL-MLR_. In setup A, we train and validate using the DeepCov dataset consisting of 3,456 proteins and test (the results here on left) using the PSICOV 150 dataset. In setup B, we train and validate using the DNCON2 dataset consisting of 1,426 proteins and test using the 62 proteins in the CASP12 dataset (the results on the right).

### Performance of DEEPCON

Finally, we compare the performance of our method DEEPCON with two state-of-the-art methods that have their training dataset publicly available. First, we compare the performance of DEEPCON with the DeepCov method when trained and tested using the same dataset consisting of 3,456 proteins. Using the covariance features as input and with the first 130 proteins as the validation set (as done in the DeepCov) we train our models and test them on the independent PSICOV dataset consisting of 150 proteins. Evaluating our predictions for the PSICOV150 dataset, we find that our implementation performs better than the DeepCov method when we develop an architecture that is similar to the one used in the DeepCov method (see P_TOP-L/5-LR_ and P_TOP-L-LR_ in **Table 1**). This could be either because of the ‘batch normalization’ layers we add and/or because of our better implementation of the overall pipeline. Next, we compare our DEEPCON method’s performance against with DeepCov method using P_TOP-L/5-LR_ and P_TOP-L-LR_. Our comparison summarized in Table 999, shows that DEEPCON achieves 5.9% improvement in P_TOP-L/5-LR_ and 15% improvement in P_TOP-L-LR_ (see **Table 1** for summary). For a fair comparison (and computing resource limitations) we only train our models once on the DeepCov dataset. In fact, to reduce our training time (and because of resource limitations), we trim all proteins to 256 residues length, possibly using lesser input information than the information used by the DeepCov method. Next, we compare DEEPCON’s performance with the DNCON2 method when trained and tested on the 1,426 proteins dataset that was used to train DNCON2. While DNCON2 uses two-level approach exploiting information from distance thresholds other than the standard 8 Angstroms and use model ensembling, we train a single model. For training our models we use the same set of 196 proteins as validation set and the remaining as the training set (as done in the DEEPCON work) and evaluate on the CASP12 dataset consisting of 84 domains. For all our predictions, we use the exact same input features (generated for targets instead of domains) as used by DNCON2. On these 84 CASP12 domains, we find that DEEPCON achieves 3.2% higher P_TOP-L/2-LR_ and 4.8% higher P_TOP-L-LR_ (see **Table 1** for details).

**Table 1.**
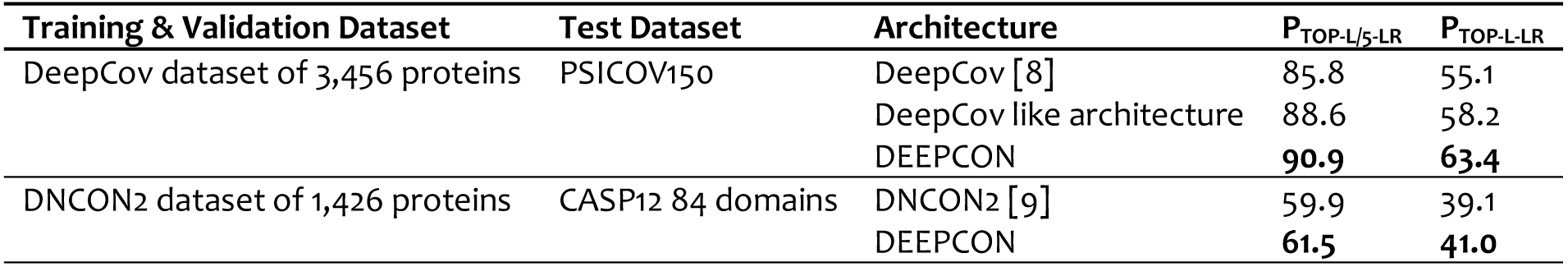
Comparison of DEEPCON’s performance with DeepCov and DNCON2.

| Training & Validation Dataset | Test Dataset | Architecture | $P_{TOP-L/5-LR}$ | $P_{TOP-L-LR}$ |
| --- | --- | --- | --- | --- |
| DeepCov dataset of 3,456 proteins | PSICOV150 | DeepCov [8] | 85.8 | 55.1 |
|  |  | DeepCov like architecture | 88.6 | 58.2 |
|  |  | DEEPCON | **90.9** | **63.4** |
| DNCON2 dataset of 1,426 proteins | CASP12 84 domains | DNCON2 [9] | 59.9 | 39.1 |
|  |  | DEEPCON | **61.5** | **41.0** |

### Evaluation on CASP 13 Dataset

Finally, we compare our DEEPCON method trained using the covariance features from the DeepCov dataset with a few state-of-the-art methods that accept multiple sequence alignment as input. For this comparison, we consider the 20 proteins targets (consisting of 32 domains) in the CASP 13 dataset for which the native structures are publicly available. Using the publicly available DNCON2 scripts for generating multiple sequence alignments, we first predicted multiple sequence alignments for these 20 protein targets. As done in the DNCON2 method, we use the sequence databases curated in 2012. Next, we predict contacts using the DEEPCON method we developed, and evaluated the contacts against the 32 domains. We also download the PconsC4 [2] tool available at https://github.com/ElofssonLab/PconsC4 and predicted contacts. Similarly, we also download the DeepCov [8] method available at https://github.com/psipred/DeepCov and predict contacts with the same alignment files as input. To obtain a baseline we also predicted contacts using CCMpred [21] and FreeContact [22]. To obtain reference results, we evaluated the contact predictions of the top group in CASP13 over these 20 targets, and also submitted the alignments to the RaptorX webserver at http://raptorx.uchicago.edu/ContactMap/ and evaluated the contacts predicted by the webserver. **Table 2** shows that our method DEEPCON performs better than the baseline methods CCMpred and FreeContact, and better then PconsC4 and DeepCov. Our comparison of DEEPCON, DeepCov, and PconsC4 with the RaptorX web-server is only for a reference because it uses many additional features other than the covariance calculated from raw alignment files. The large difference between the best group’s performance in CASP13 and the RaptorX webserver (in **Table 2**) highlights the performance gain that could be achieved with better/larger multiple sequence alignments. This is because the top group in CASP13 experiment was the RaptorX method itself. The differences of precision between the baseline methods (CCMpred and FreeContact) and the deep learning methods (DEEPCON, DeepCov, and PconsC4) represents the gain from the use of deep neural networks.

**Table 2.** Comparison of DEEPCON’s performance with other methods that accept multiple sequence alignment as input on the 32 domains in the CASP13 dataset of 20 protein targets. Methods with an asterisk (*) are listed for reference.

| | $P_{TOP-L/5-LR}$ | $P_{TOP-L-LR}$ | $P_{TOP-L/5-MLR}$ | $P_{TOP-L-MLR}$ |
| --- | --- | --- | --- | --- |
| * Best Group in CASP13 | 79.6 | 55.2 | 88.8 | 66.5 |
| * RaptorX Webserver | 67.4 | 43.8 | 75.3 | 53.2 |
| DEEPCON (our method) | 57.2 | 34.2 | 67.5 | 45.9 |
| DeepCov [8] | 48.2 | 28.3 | 61.6 | 37.9 |
| PconsC4 [2] | 42.7 | 27.3 | 53.1 | 34.9 |
| CCMpred [21] | 32.8 | 19.3 | 34.8 | 22.6 |
| FreeContact [22] | 30.3 | 17.4 | 33.5 | 20.0 |

### Conclusions

We found that regularization using dropout is highly effective in all architectures - fully convolutional, residual, or dilated residual networks. We also found that dilated convolutional methods yield negligibly better performance than the regular residual networks, but these gains are amplified when dilated convolutions are combined with dropouts. We believe that our findings when combined with techniques such as using multiple distance thresholds [9,15], ensembling [3,9], predicting actual distances using binning [17], etc. used in other methods can significantly improve the overall contact prediction precision. We also believe that our findings and the architectures will be utilized by other researchers to continue the development and obtain better performance with more complex and powerful architectures, larger training and validation datasets, development of additional features and output labels, and through model ensembling.

## Acknowledgements

Some of the computation for this work was performed on the high-performance computing infrastructure provided by Research Computing Support Services (RCSS) and in part by the National Science Foundation under grant number CNS-1429294 at the University of Missouri, Columbia MO. We would like to thank the RCSS team for their infrastructure and technical support. We also thank Google Cloud for cloud credits through the GCP cloud credits program. We also gratefully acknowledge the support of NVIDIA Corporation with the donation of the Quadro P6000 GPU used for this some of the computations in this research.

## References

1. Berman HM. The Protein Data Bank. Nucleic Acids Res. 2000;28: 235–242. doi:10.1093/nar/28.1.235

2. Michel M, Hurtado DM, Elofsson A. PconsC4: fast, free, easy, and accurate contact predictions. bioRxiv. Cold Spring Harbor Laboratory; 2018; 383133. doi:10.1101/383133

3. Wang S, Sun S, Li Z, Zhang R, Xu JJJJ, Trakselis M, et al. Accurate De Novo Prediction of Protein Contact Map by Ultra-Deep Learning Model. Schlessinger A, editor. PLOS Comput Biol. ICML; 2017;13: e1005324. doi:10.1371/journal.pcbi.1005324

4. Moult J, Fidelis K, Kryshtafovych A, Schwede T, Tramontano A. Critical assessment of methods of protein structure prediction (CASP)-Round XII. Proteins Struct Funct Bioinforma. 2018;86: 7–15. doi:10.1002/prot.25415

5. Monastyrskyy B, D’Andrea D, Fidelis K, Tramontano A, Kryshtafovych A. New encouraging developments in contact prediction: Assessment of the CASP11 results. Proteins Struct Funct Bioinforma. 2016;84: 131–144. doi:10.1002/prot.24943

6. Mirdita M, Von Den Driesch L, Galiez C, Martin MJ, Soding J, Steinegger M. Uniclust databases of clustered and deeply annotated protein sequences and alignments. Nucleic Acids Res. 2017; doi:10.1093/nar/gkw1081

7. Suzek BE, Huang H, McGarvey P, Mazumder R, Wu CH. UniRef: Comprehensive and non-redundant UniProt reference clusters. Bioinformatics. 2007; doi:10.1093/bioinformatics/btm098

8. Jones DT, Kandathil SM. High precision in protein contact prediction using fully convolutional neural networks and minimal sequence features. Valencia A, editor. Bioinformatics. 2018; doi:10.1093/bioinformatics/bty341

9. Adhikari B, Hou J, Cheng J. DNCON2: improved protein contact prediction using two-level deep convolutional neural networks. Bioinformatics. 2017; doi:10.1093/bioinformatics/btx781

10. Mirabello C, Wallner B. rawMSA: proper Deep Learning makes protein sequence profiles and feature extraction obsolete. doi:10.1101/394437

11. Kosciolek T, Jones DT. Accurate contact predictions using coevolution techniques and machine learning. Proteins Struct Funct Bioinforma. 2015; n/a-n/a. doi:10.1002/prot.24863

12. Michel M, Skwark MJ, Menéndez Hurtado D, Ekeberg M, Elofsson A. Predicting accurate contacts in thousands of Pfam domain families using PconsC3. Bioinformatics. 2017;33: 2859–2866. doi:10.1093/bioinformatics/btx332

13. Eigen D, Puhrsch C, Fergus R. Depth Map Prediction from a Single Image using a Multi-Scale Deep Network.

14. Wang G, Dunbrack RL. PISCES: A protein sequence culling server. Bioinformatics. 2003; doi:10.1093/bioinformatics/btg224

15. Jones DT, Singh T, Kosciolek T, Tetchner S. MetaPSICOV: Combining coevolution methods for accurate prediction of contacts and long range hydrogen bonding in proteins. Bioinformatics. 2014;31: btu791. doi:10.1093/bioinformatics/btu791

16. Ronneberger O, Fischer P, Brox T. U-net: Convolutional networks for biomedical image segmentation. Lecture Notes in Computer Science (including subseries Lecture Notes in Artificial Intelligence and Lecture Notes in Bioinformatics). 2015. pp. 234–241. doi:10.1007/978-3-319-24574-4_28

17. Xu J. Distance-based Protein Folding Powered by Deep Learning. bioRxiv. Cold Spring Harbor Laboratory; 2018; 465955. doi:10.1101/465955

18. Söding J. Protein homology detection by HMM–HMM comparison. Bioinformatics. 2005;21: 951–60. doi:10.1093/bioinformatics/bti125

19. Finn RD, Clements J, Eddy SR. HMMER web server: interactive sequence similarity searching. Nucleic Acids Res. 2011;39: W29–W37. doi:10.1093/nar/gkr367

20. Zagoruyko S, Komodakis N. Wide Residual Networks. 2016;

21. Seemayer S, Gruber M, Söding J. CCMpred - Fast and precise prediction of protein residue-residue contacts from correlated mutations. Bioinformatics. 2014;30: 3128–3130. doi:10.1093/bioinformatics/btu500

22. Kaján L, Hopf T a, Kalaš M, Marks DS, Rost B. FreeContact: fast and free software for protein contact prediction from residue co-evolution. BMC Bioinformatics. 2014;15: 85. doi:10.1186/1471-2105-15-85

